# Shaken, not shifted: Genotypic variation tunes how interspecific competition shapes niches

**DOI:** 10.1101/2023.03.15.532847

**Authors:** Raul Costa-Pereira, Inês Fragata, André Mira, Maud Charlery de la Masselière, Joaquin Hortal, Sara Magalhães

## Abstract

Individual variation in resource use as well as in the response to competitors has been recognized as playing an important role is species interactions. Still, we have as yet little information on whether such responses have a genetic basis as well as on how they affect each other. Here, we tested whether 20 genetically inbred lines of the spider mite *Tetranychus evansi* vary in their response to a gradient of cadmium concentration within plants as well as in their propensity to reshape niches when facing interspecific competition along this gradient. In absence of interspecific competitors, most lines were negatively affected by cadmium, albeit often in a non-linear fashion. Morevoer, half of the lines exhibited changes in the curvature of the relationship between number of females and cadmium concentration when facing competition with the congeneric *T. urticae*. Inbred lines also showed a shallower decay in offspring number along the cadmium gradient in presence of interspecific competition. Our findings provide evidence for large, partly genetic, variation in resource use and in the response to interspecific competition in heterogeneous environments. Moreover, we show that genotype responses to interspecific competition is contingent upon their response to an environmental gradient. Together, our results thus emphasize the importance of considering intraspecific variation in responses to interspecific competition, providing novel insights to link intra- and interspecific levels of biodiversity.

## Introduction

Interspecific competition is a powerful interaction shaping the ecology and evolution of interacting species. Throughout the “competitive” history of ecology (Raerinne 2020), ecologists have nurtured particular interest in documenting how co-occurring species respond to competition for limited resources (Paine 1966). Numerous experimental and observational studies show that competition with heterospecifics leads to changes in niches, life history, and spatio-temporal distribution of organisms across diverse scales, taxa, and ecosystems (Tilman 1982, Connell 1983, Kaplan and Denno 2007, Aschehoug et al. 2016).

The consequences of interspecific competition depend on both intrinsic and extrinsic factors. Although this is a not novel realization for ecologists (Schoener 1983, Thompson 1988), we still lack a comprehensive understanding on the interplay between factors related to the interacting organisms per se (i.e., intrinsic) and to the environmental context in which interactions take place (i.e., extrinsic). Classic theories have approached the extrinsic component by making predictions about how habitat properties (e.g., abiotic gradients, environmental stability) affect species’ responses to competition (Roughgarden 1983, Tilman 2020). For instance, habitat heterogeneity offers scope for competing species to find competitive refugia and partition spatially their niches (Levin 1976, Danielson 1991). However, competitors often present similar niche preferences, which creates a tension between the detrimental consequences of competition and the beneficial effects of optimal resource or habitat choice (Pimm and Rosenzweig 1981, Abrams 1987). Although this duality has been relatively well-studied from a theoretical perspective and for specific taxa, we still know little whether and how intrinsic factors play a role mediating how competition shapes niches in heterogeneous environments.

Considering species-level traits is the classical approach to test the effect of competition on species distribution (Godoy et al. 2014, Kraft et al. 2015). However, although phenotypic diversity is more conspicuous across species than within them, selective pressures affect trait selection at the individual level (Roughgarden 1972). Due to this, variation within and across populations of the same species can play a major role on competition dynamics (Gómez-Mestre and Tejedo 2002, Violle et al. 2012, Des Roches et al. 2018). Such intraspecific ecological diversity can emerge through multiple, non-mutually exclusive ecological and evolutionary mechanisms, but seldom ecological studies consider the origin of observed variation within species (Bolnick et al. 2003, Dall et al. 2012, Costa-Pereira and Araújo 2022). In particular, genotypic diversity across populations is a potential source of variation in competitive responses (Cahill Jr et al. 2005), but it has been underexplored empirically. The geographic mosaic of selective forces has the potential to create and maintain variation across populations regarding how species deal with biotic and abiotic challenges (Travis 1996, Post et al. 2008). Understanding whether intraspecific variation leads to different competitive responses across populations can contribute, for example, to develop more accurate predictions on interactions outcomes across the geographic space (Leisnham et al. 2009, Moran et al. 2016). In this sense, studying how interspecific competition shapes niches across multiple genetic lines, and its cascading consequences to performance, has the potential to shed new light on the interplay between extrinsic and intrinsic factors driving competitive outcomes.

Here, we investigated how intraspecific variation tunes the effects of interspecific competition on habitat selection and reproductive performance along an environmental gradient, using individuals from 20 inbred lines of the herbivorous spider mite *Tetranychus evansi*, when facing or not competition with the congeneric *T. urticae*. In this system, genetic lines can be used as a proxy for different individuals (Godinho et al. 2020a), so their study provides insights on how ecological responses vary between conspecific individuals of the same species. These two spider mite species co-occur in natural communities (Ferragut et al. 2013) and compete for common resources on tomato plants (Godinho et al. 2020b, Fragata et al. 2022). Tomato plants accumulate cadmium from the environment, which has cascading detrimental effects on spider mites’ distribution and fitness (Godinho et al. 2018, Godinho et al. 2022). Natural populations of both spider mite species show variation in their preference and performance across a cadmium gradient (Godinho et al. 2022). Therefore, we expect that interspecific competition displaces *T. evansi* from low-cadmium habitat patches, shifting their realized niches towards sub-optimal high-cadmium patches. If this shift in the niche happens as we expect, exploring resources from high-cadmium patches should result in reduced offspring numbers. Because competition and cadmium-mediated stress are powerful selection pressures in this system (Godinho et al. 2022), we expect genetic lines to present similar responses to these biotic and abiotic factors.

Specifically, we first predict that (1) females of *T. evansi* prefer low-cadmium plants. Then, we predict that the presence of *T. urticae* in low-cadmium habitats (2) increases the occupancy of high-cadmium plants and ultimately (3) reduces offspring number in *T. evansi*. To contrast preference and reproductive fitness patterns when facing competition, we quantified metrics that represent how spider mites select where to forage and to oviposit along the cadmium gradient, which allows us to quantify, respectively, whether multiple genetic lines reshape niches and change offspring due to interspecific competition.

## Methods

### Plant and spider mite populations

Tomato plants (*Solanum lycopersicum*, var MoneyMaker) used in the experiment were sown in a climatic room (25 ºC, 64% of humidity, light:dark = 16:8). Tomato seeds were watered twice per week until the plants reached two-week-old, where four Cd chloride solution with different concentrations (0mM, 0.5 mM, 1.5 mM and 3mM) were added to the watering process. After five weeks, plants were used in the experiments and had at least five fully expanded leaves.

*Tetranychus evansi* was used as the focal species, and *Tetranychus urticae* as the competitor. For each species, an outbred population was created through controlled crossings of populations collected from field tomato plants, in Portugal in 2017 (Zélé et al. 2018, Godinho et al. 2020a). Mites were maintained on detached tomato leaves placed in isolated boxes under controlled conditions (23.5°C, 68% humidity, light:dark = 16:8) in a climate-room at the University of Lisbon. The stems of these leaves were inside a small container with water. Twice a week, water and new tomato leaves were added, and overexploited leaves were removed.

From the *T. evansi* outbred population, we created 100 inbred lines from 14 generations of sib mating, thus ensuring an inbreeding coefficient of more than 96% (Godinho et al. 2020a). Each line thus roughly corresponds to one genotype. Twenty of these lines were used in the current experiment. All lines were maintained on small leaf discs inside Petri dishes. Every 14 days, 25 adult females of each line were transferred to a new prepared Petri dish. To ensure that females used in the experiments were approximately of the same age, adult females of each *T. evansi* isogenic lines and of the *T. urticae* population where isolated on separate tomato leaves and allowed to lay eggs for 48 h. In all experiments, we used adult females resulting from this cohort, 15 days after egg hatching.

### Experimental setup

To experimentally create environmental heterogeneity for spider mites, we used the sown tomato plants to conceive an artificial habitat with increasing cadmium concentrations: 0, 0.5, 1.5 and 3 mM. One plant per concentration was used, and all were connected by a platform that allowed the movement of the mites between plants. This platform was connected to the 4th leaf of each plant, itself isolated from the rest of the plant via a lanoline barrier applied to its stems. In this way, mites could only occur on that leaf, which facilitated the subsequent counting. Two treatments were established, with or without interspecific competitors. The former was generation by introducing 100 females of *T. urticae* to tomato plants grown with 0 and 0.5 mM of cadmium (corresponding to the favorable part of the gradient) 24 hours prior to the addition of *T. evansi* females. To both treatments, 200 females of *T. evansi* were placed in the platform to allow plant selection. Eight replicates per treatment were performed.

### Quantifying habitat selection and offspring performance

To quantify niche shape in a context of habitat choice, 48h after placing the females, we counted the number of *T. evansi* present on the 4^th^ leaf of each plant. Subsequently, to assess the offspring performance, the 4^th^ leaf was isolated, and all adult females were removed, leaving only the eggs. New leaves were added to provide enough resources for the development of the offspring. The number of female offspring was counted 15 days later.

### Effects of interspecific competition on habitat selection across lines

For each genetic line of *T. evansi*, we tested if (i) habitat selection patterns and (ii) offspring number along the cadmium gradient change when interspecific competition with was present or absent. We fit generalized linear mixed-effects models with (i) the number of females that choose each habitat patch (i.e., a plant with a given cadmium concentration) or (ii) the number of offspring per female as response variable, and cadmium concentration and competition as predictors. To control for the data when the experiment was performed, we included the experimental block as random factor. Specifically, because competitive responses are not necessarily linear, we fit a linear and polynomial up to 3^rd^ degree and then estimated the Akaike information criterion (AIC) for each model. For both response variables, most lines showed as best fit (i.e., lowest AIC) the polynomial model, and no line presented the best model that differed from the mode (i.e., a difference between models < −2). Thus, for the sake of simplicity, we used the polynomial model for all lines. In this context, we used the estimated leading coefficients (x^2^) to determine how competitors change niche shape for each genetic line. Specifically, if the leading coefficients are lower than 0 then the choice is concentrated to intermediate cadmium concentrations (or at least not on 0), if the leading coefficients are higher than 0 then lower choice is in middle to higher cadmium concentrations.

As an alternative approach to quantify habitat choice along the cadmium gradient when predators are present or absent, we re-ran linear mixed models using the log-transformed number of females in each habitat and then extracted the resulting intercepts and slopes. In this approach, resulting slopes can be used as a metric to quantify the strength of preference towards low-cadmium plants. For example, highly negative slopes indicate a sharp decrease in choice towards increased cadmium concentrations.

## Results

In the absence of interspecific competition, *T. evansi* females predominantly choose plants with low to moderate cadmium concentrations. Overall, this preference is consistent across the genetic lines (Figures 1, S1), revealing intraspecific niche conservatism. However, some of lines presented weak (e.g., line #9) or even opposing (e.g., line #23) preference patterns (Figure S1, S2).

**Figure 1.**
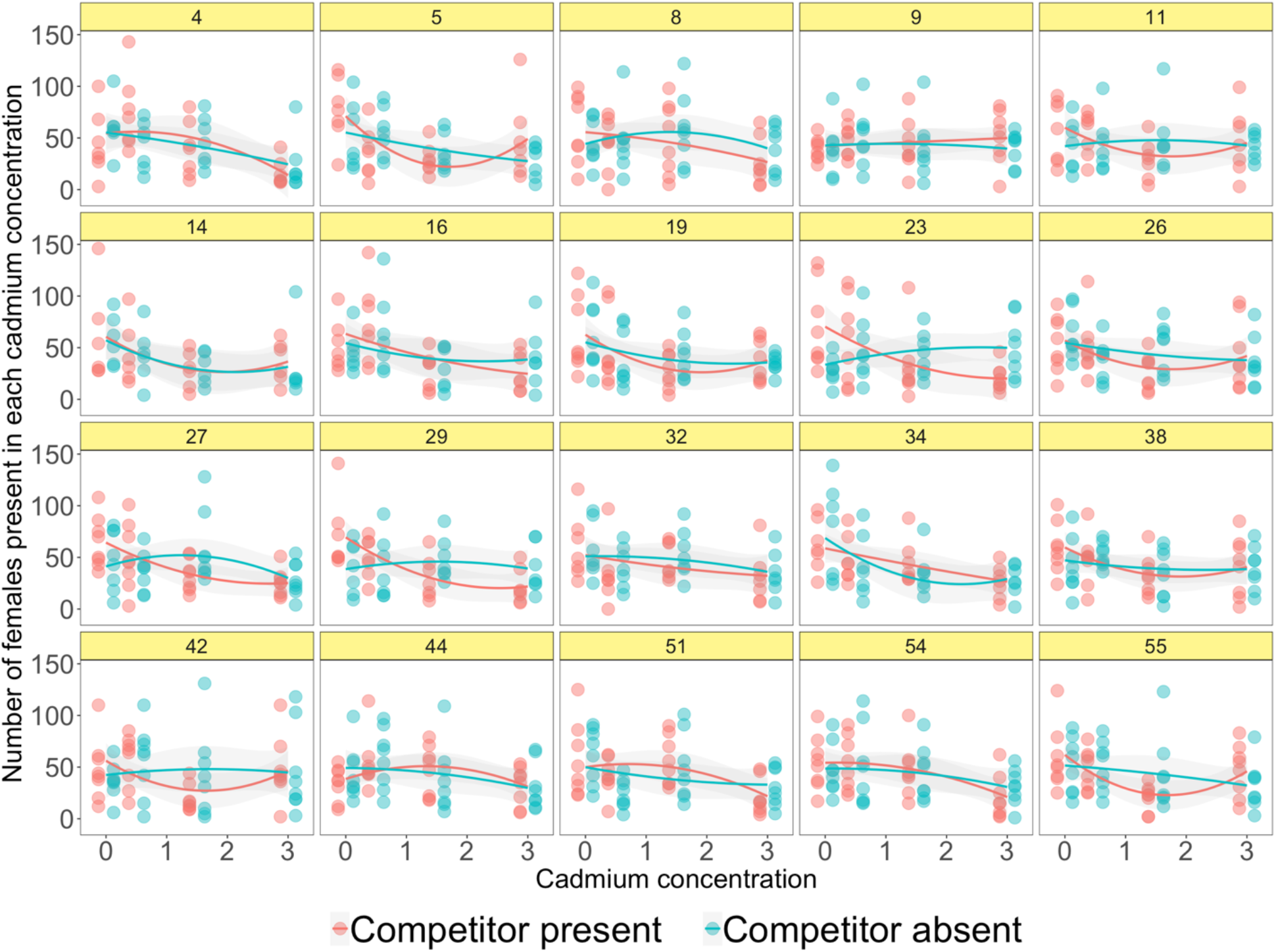
Number of *Tetranychus evansi* females that selected plants along a gradient of four cadmium concentrations (0, 0.5, 1.5 and 3 mM) when heterospecific competitors (*T. urticae*) are present or absent. the treatment in which competitors were present was created by adding females of the congeneric *T. urticae* to the plants with 0 and 0.5 mM of cadmium 24h before *T. evansi* females. Each panel represents the responses of one of 20 different genetic lines.

At first sight, the presence of the competitor *T. urticae* on low-cadmium plants had just a modest effect on the habitat choice by *T. evansi* females (Figure 1) as low and moderate cadmium concentrations remain, in general, preferred despite the presence of competitors (Figures 1, S1). However, there is relevant variation in the shape of this response across genetic lines. Indeed, in half of the lines (n = 10), interspecific competition changed the curvature of the relationship between number of females and cadmium concentration, while the other half kept the same shape of response (Figure 2; Table S1). For the 10 lines that exhibited changes in curvature, most (n = 8) went from a downward (concave) shape in absence of competition (i.e., lower habitat occupancy in medium-cadmium concentration) to an upward (convex) shape when facing interspecific competition (i.e., higher habitat occupancy in medium-cadmium plants) (Figure 2). Slopes from the models using log-transformed number of females as response variable ratify this pattern by showing that while for most of the lines the presence of the competitor increases occupancy towards lower-cadmium concentrations, some lines presented no signs of preference at all or even an increased choice towards high-cadmium plants (Figure S2).

**Figure 2.**
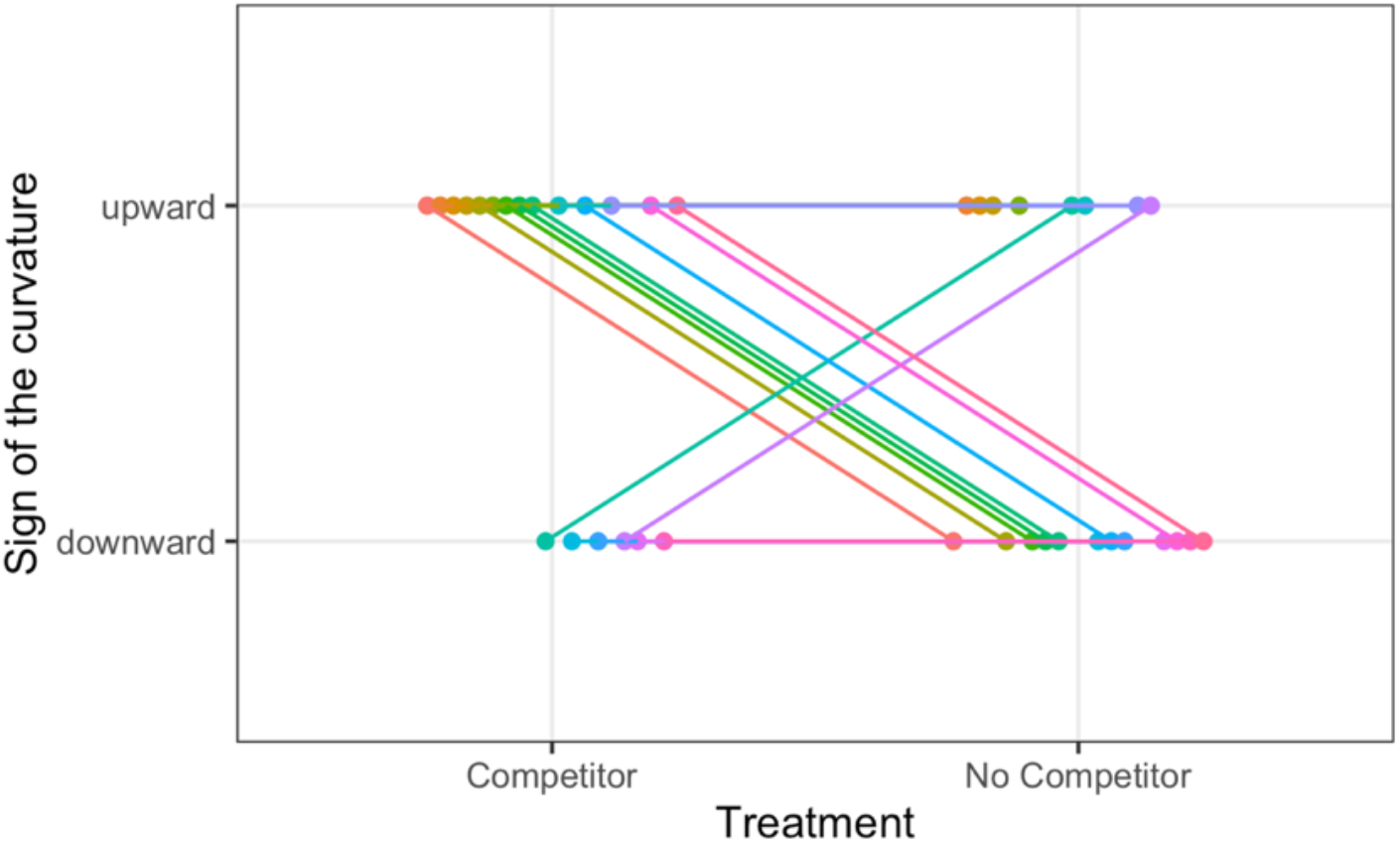
Polynomial models to quantify how interspecific competition with *Tetranychus urticae* changes habitat selection by females of *T. evansi* along a gradient of cadmium concentration. This plot shows how the curvature of resulting models respond (i.e., reshape to downward, reshape to upward or no change in curvature) across 20 genetic lines (colors represent different genetic lines). Downward curvatures mean a lower habitat selection in medium-cadmium concentration, while upward curvatures mean a higher habitat selection in medium-cadmium plants. Half of the genetic lines did not change the curvature in response to interspecific competition (represented by horizontal lines: four lines kept downward and six did so with upward curvatures).

We observed less variation across genetic lines in how offspring numbers respond to cadmium concentration and competition (Figure 3, Table S2). When interspecific competitors where absent, offspring number decayed sharply at higher cadmium concentrations, resulting in a general upward (convex) response (Figure 3). Overall, interspecific competition led to a decrease in offspring number, but many lines presented a slight increase in their offspring number in intermediate cadmium concentrations, coupled with a decrease in the number of offspring present in the lower cadmium concentrations (Figure 3). Interestingly, just six genetic lines changed the curvature to a downward shape when competitors where present (Figure 4, Table S2), suggesting that interspecific competition does not affect the proportion of offspring on the different cadmium concentrations for most lines. Just two of these six lines (lines #34 and #51) also exhibited changes in the curvature related to habitat choice (i.e., niche shape) in response to interspecific competition (Table S1 and S2).

**Figure 3.**
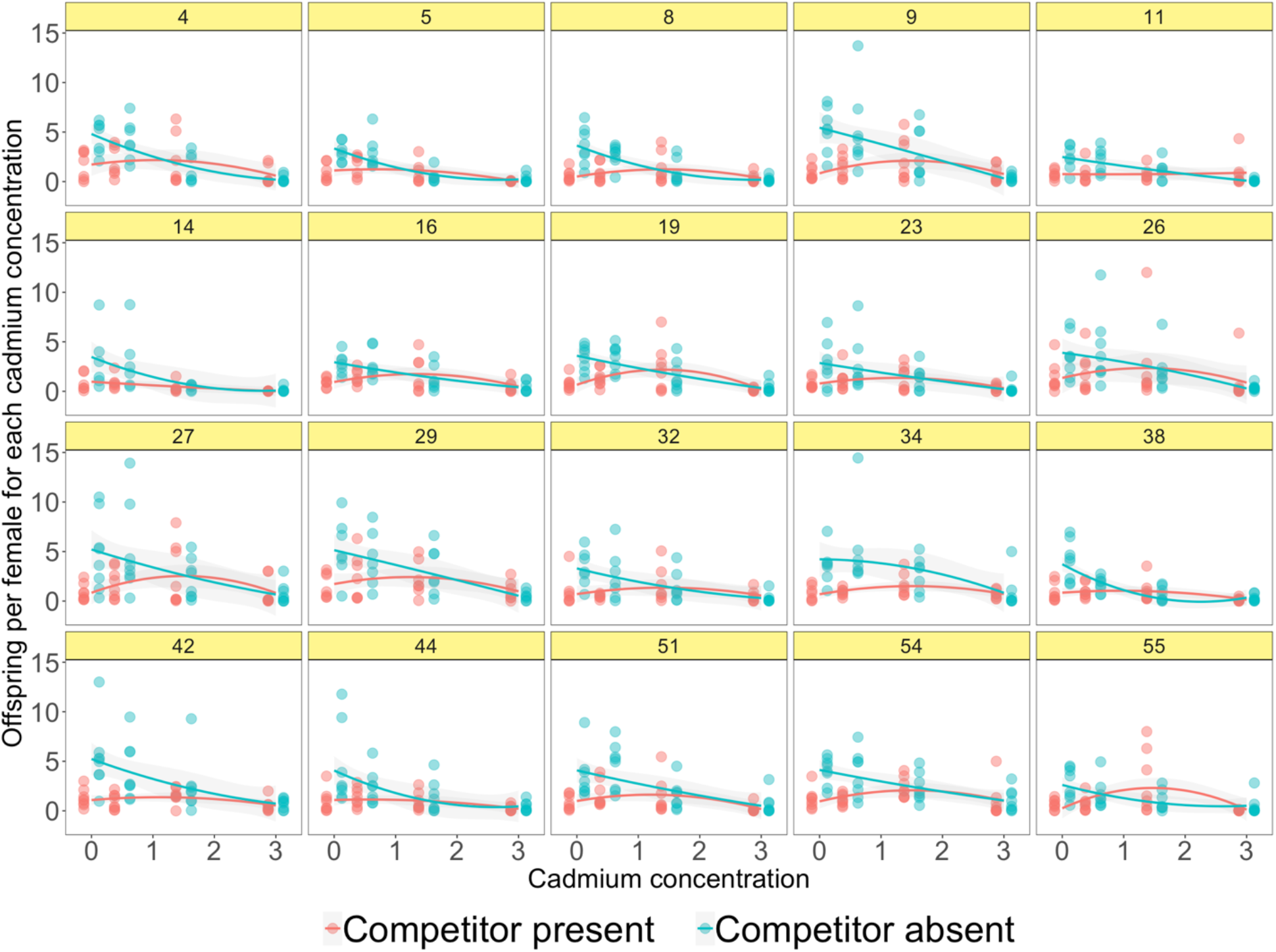
Offspring per female of *Tetranychus evansi* along a gradient of four cadmium concentrations (0, 0.5, 1.5 and 3 mM) across 20 different genetic lines. Each panel represents the responses of one of 20 different genetic lines.

**Figure 4.**
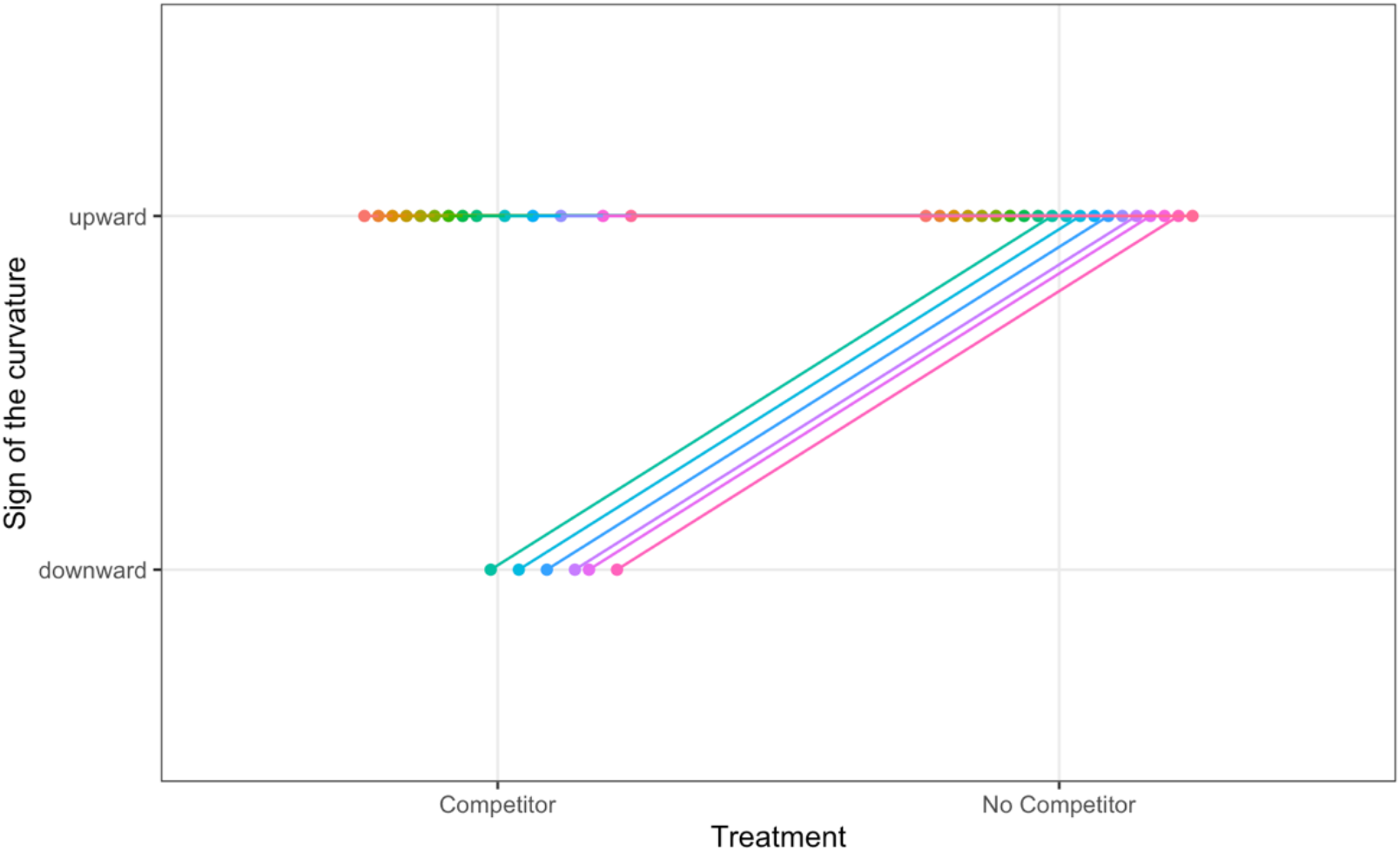
Responses of the signs of the curvature from polynomial models to quantify how interspecific competition with *Tetranychus urticae* changes offspring number by females of *T. evansi* along a gradient of cadmium concentration. All lines presented upward curvatures when competitors were absent and just six lines changes the curvature to downward when competitors were present.

## Discussion

Ecologists have long recognized that interspecific competition can reshape niches and reduce reproductive fitness across different taxa (Hairston et al. 1960). However, the generality of these effects in heterogeneous habitats and across genetic lines within interacting species remains uncertain. Our experiments on the effects of interspecific competition on habitat choice and offspring number show that the females of the spider mite *T. evansi*. prefer low-cadmium plants, even when the heterospecific competitor *T. urticae* occupies these preferred habitats. Despite the prevalence of these responses across populations, some genetic lines present specific responses to both cadmium and competition. Interestingly, heterospecific competitors reshaped (by changing the curvature) the habitat selection curve in half of the studied lines, in general favoring an increase in the choice towards medium-cadmium concentrations. Finally, we found a general decay in offspring number as cadmium concentration increases, but interspecific competition changed the shape of this pattern in several genetic lines. Together, these results highlight how interspecific interactions can trigger diverse responses across genetic lines within species, which has important implications to how we quantify and predict interactions in a rapidly changing world.

Perhaps the most striking of these results is that we observed no sign of a general niche response to competition across genetic lines. Half of the studied lines did not exhibit substantial shifts in the niche (i.e., proportion of habitat selected along the cadmium gradient) in response to competition. For these lines, it is possible that the beneficial consequences of exploring no-or low-cadmium plants may overcome the detrimental impacts of interspecific competition. In turn, the other 10 lines presented two distinct responses: either increase or decrease selection of medium-cadmium concentrations. Although we cannot pinpoint the underlying ecological and evolutionary mechanisms driving this variation across genetic lines, our findings reveal that intraspecific variation in response to competition is a potential mechanism influencing species distribution and abundance along environmental gradients. In this sense, even under the very same extrinsic context (i.e., same abiotic gradient and same density of competitors), different populations may differ in their niches and spatial arrangement due to intrinsic factors.

This possibility challenges the classic assumption that species realized niches emerge from the projection of fundamental niche dimensions into particular combinations of extrinsic factors, namely resources and conditions (Hutchinson 1978, Colwell and Rangel 2009). Under this perspective, intraspecific variation in how species respond to biotic interactions (such as competition) questions the widely accepted idea that the occurrence of species in local communities are driven by the mere overlap of species responses to abiotic and biotic gradients, mediated by dispersal processes (Soberón 2007, Soberón and Nakamura 2009, Hortal et al. 2010). Rather, it definitively calls for accounting for individual responses to the complex combinations of these gradients, as advocated by Roughgarden (1972). This individualistic perspective is common in disciplines such as evolutionary ecology and (plant) functional ecology, but not so in many approaches to community ecology.

These intraspecific variations are not the major driver of *T. evansi* reproductive responses to the abiotic gradient we studied, though. The decay in offspring numbers as cadmium concentration increases was similar across genetic lines. Reproductive performance under interspecific competition were also not as variable across genetic lines as niche responses (i.e., habitat selection patterns). This suggests that different niche shapes across lines can result in similar fitness outcomes in heterogeneous habitats. Another convergent pattern across lines is the overall lower offspring number when *T. urticae* was present, in line with expectations from competition theory and studies from diverse taxa (Gurevitch et al. 1992, Kaplan and Denno 2007). These convergent responses across lines confirm that interspecific competition is indeed a powerful driver of spider mite’s performance (Godinho et al. 2020b, Fragata et al. 2022).

Interestingly, when interspecific competitors are present, most lines showed a slight increase in the offspring number in intermediate cadmium concentrations. Accordingly, there was a prevalent decrease in offspring number in low cadmium concentrations. Together, these results suggests that suboptimal habitat patches when competitors are absent may become good choices to improve performance when competitors arrive first in these preferred habitats, which has also implications for coexistence in spider mites (Fragata et al. 2022).

Elucidating which local biotic and abiotic contexts have driven observed variation across genetic lines remains an open question. Indeed, *T. urticae* and *T. evansi* co-occur in natural communities (Ferragut et al. 2013), but the magnitude of competition is likely to vary geographically depending on, for instance, local relative abundances and resource availability (see Leibold et al. 2019). Also, cadmium concentrations in soil vary largely across sites (Kubier et al. 2019). Therefore, a geographic mosaic of abiotic (cadmium concentration) and biotic (competition intensity) gradients has the potential to generate and maintain intraspecific diversity in responses to competition across populations. An important next step in this direction is to clarify how the observed contrasting responses to competition affect the coexistence of *T. urticae* and *T. evansi*. Competitive outcomes in this system are not trivial, depending, for instance, on the order of species arrival (Fragata et al. 2022). Our results add a new layer of complexity to this scenario by revealing the plastic niche responses of *T. evansi* to competition. In this sense, future studies should address whether and how competitive outcomes (e.g., coexistence or competitive exclusion) in heterogeneous environments vary across genetic lines.

To summarize, our experiments demonstrate that species can show diverse intraspecific responses to interspecific competition in heterogeneous environments. The observed variation across genetic lines in the propensity to reshape niches under interspecific competition emphasizes that models predicting the outcome of species interactions should not neglect intraspecific variance in how species deal with competition. Such intraspecific, individual variability may impact key aspects of species responses to changing environments, determining interaction strengths (e.g., Costa-Pereira et al. 2018, Costa-Pereira et al. 2019) or the establishment of populations under novel habitat conditions (Tamme et al. 2010, Mestre et al. 2020). This is particularly important for the generation of diversity within regional communities, as these within-species variations may help accommodating higher numbers of species occupying overlapping niche space (Ricklefs 2010). This has broad implications for when predicting the outcome of interactions across large geographic areas, as intraspecific variation in responses to competition may be amplified as species experience distinct environmental contexts (Moran et al. 2016). Consequently, as biotic interactions and abiotic conditions have been rapidly changing in diverse ecosystems worldwide, intraspecific variation in species responses to competition (and other interspecific interactions) may boost spatial variation in interaction outcomes (Arroyo-Correa et al. 2023). This calls for a reinvigoration of the research in the ecological differences between individuals (Dall et al. 2012). In sum, although this lack of generality in competitive responses challenges key assumption of current competition models, it also opens the way for potential evolutionary responses within short time frames and provides novel insights to link intra- and interspecific levels of biodiversity.

## Acknowledgments

This work was funded by ERC (European Research Council) consolidator grant COMPCON, GA 725419 led by SM, and by FCT (Fundação para Ciência e Tecnologia) with the Junior researcher contract (CEECIND/02616/2018) attributed to IF. RC-P is supported by the Serrapilheira Institute (grant number Serra – R-2011-37572) and grant #2020/11953-2, São Paulo Research Foundation (FAPESP).

## References

Abrams, P. A. 1987. Alternative models of character displacement and niche shift. I. Adaptive shifts in resource use when there is competition for nutritionally nonsubstitutable resources. Evolution 41:651–661.

Arroyo-Correa, B., P. Jordano, and I. Bartomeus. 2023. Intraspecific variation in species interactions promotes the feasibility of mutualistic assemblages. Ecology Letters 26:448–459.

Aschehoug, E. T., R. Brooker, D. Z. Atwater, J. L. Maron, and R. M. Callaway. 2016. The Mechanisms and Consequences of Interspecific Competition Among Plants. Annual Review of Ecology, Evolution, and Systematics 47:263–281.

Bolnick, D. I., R. Svanbäck, J. A. Fordyce, L. H. Yang, J. M. Davis, C. D. Hulsey, and M. L. Forister. 2003. The ecology of individuals: incidence and implications of individual specialization. The American Naturalist 161:1–28.

Cahill Jr, J. F., S. W. Kembel, and D. J. Gustafson. 2005. Differential genetic influences on competitive effect and response in Arabidopsis thaliana. Journal of Ecology 93:958–967.

Colwell, R. K., and T. F. Rangel. 2009. Hutchinson’s duality: The once and future niche. Proceedings of the National Academy of Sciences 106:19651–19658.

Connell, J. H. 1983. On the Prevalence and Relative Importance of Interspecific Competition: Evidence from Field Experiments. The American Naturalist 122:661–696.

Costa-Pereira, R., and M. S. Araújo. 2022. Individual Specialization.in S. Scheiner, editor. Encyclopedia of Biodiversity. Elsevier.

Costa-Pereira, R., M. S. Araújo, F. L. Souza, and T. Ingram. 2019. Competition and resource breadth shape niche variation and overlap in multiple trophic dimensions. Proceedings of the Royal Society B: Biological Sciences 286:20190369.

Costa-Pereira, R., V. H. W. Rudolf, F. L. Souza, and M. S. Araújo. 2018. Drivers of individual niche variation in coexisting species. Journal of Animal Ecology 87:1452–1464.

Dall, S. R. X., A. M. Bell, D. I. Bolnick, and F. L. W. Ratnieks. 2012. An evolutionary ecology of individual differences. Ecology Letters 15:1189–1198.

Danielson, B. J. 1991. Communities in a Landscape: The Influence of Habitat Heterogeneity on the Interactions between Species. The American Naturalist 138:1105–1120.

Des Roches, S., D. M. Post, N. E. Turley, J. K. Bailey, A. P. Hendry, M. T. Kinnison, J. A. Schweitzer, and E. P. Palkovacs. 2018. The ecological importance of intraspecific variation. Nature Ecology & Evolution 2:57–64.

Ferragut, F., E. Garzón-Luque, and A. Pekas. 2013. The invasive spider mite Tetranychus evansi (Acari: Tetranychidae) alters community composition and host-plant use of native relatives. Experimental and Applied Acarology 60:321–341.

Fragata, I., R. Costa-Pereira, M. Kozak, A. Majer, O. Godoy, and S. Magalhães. 2022. Specific sequence of arrival promotes coexistence via spatial niche pre-emption by the weak competitor. Ecology Letters 25:1629–1639.

Godinho, D. P., M. A. Cruz, M. Charlery de la Masselière, J. Teodoro-Paulo, C. Eira, I. Fragata, L. R. Rodrigues, F. Zélé, and S. Magalhães. 2020a. Creating outbred and inbred populations in haplodiploids to measure adaptive responses in the laboratory. Ecology and Evolution 10:7291–7305.

Godinho, D. P., A. Janssen, D. Li, C. Cruz, and S. Magalhães. 2020b. The distribution of herbivores between leaves matches their performance only in the absence of competitors. Ecology and Evolution 10:8405–8415.

Godinho, D. P., H. C. Serrano, A. B. Da Silva, C. Branquinho, and S. Magalhães. 2018. Effect of Cadmium Accumulation on the Performance of Plants and of Herbivores That Cope Differently With Organic Defenses. Frontiers in Plant Science 9.

Godinho, D. P., H. C. Serrano, S. Magalhães, and C. Branquinho. 2022. Concurrent herbivory and metal accumulation: The outcome for plants and herbivores. Plant-Environment Interactions 3:170–178.

Godoy, O., N. J. B. Kraft, and J. M. Levine. 2014. Phylogenetic relatedness and the determinants of competitive outcomes. Ecology Letters 17:836–844.

Gómez-Mestre, I., and M. Tejedo. 2002. Geographic variation in asymmetric competition: a case study with two larval anuran species. Ecology 83:2102–2111.

Gurevitch, J., L. L. Morrow, A. Wallace, and J. S. Walsh. 1992. A Meta-Analysis of Competition in Field Experiments. The American Naturalist 140:539–572.

Hairston, N. G., F. E. Smith, and L. B. Slobodkin. 1960. Community Structure, Population Control, and Competition. The American Naturalist 94:421–425.

Hortal, J., N. Roura-Pascual, N. J. Sanders, and C. Rahbek. 2010. Understanding (insect) species distributions across spatial scales. Ecography 33:51–53.

Hutchinson, G. E. 1978. An Introduction to Population Biology. Yale University Press, New Haven, CT.

Kaplan, I., and R. F. Denno. 2007. Interspecific interactions in phytophagous insects revisited: a quantitative assessment of competition theory. Ecology Letters 10:977–994.

Kraft, N. J. B., O. Godoy, and J. M. Levine. 2015. Plant functional traits and the multidimensional nature of species coexistence. Proceedings of the National Academy of Sciences 112:797–802.

Kubier, A., R. T. Wilkin, and T. Pichler. 2019. Cadmium in soils and groundwater: A review. Applied Geochemistry 108:104388.

Leibold, M. A., M. C. Urban, L. De Meester, C. A. Klausmeier, and J. Vanoverbeke. 2019. Regional neutrality evolves through local adaptive niche evolution. Proceedings of the National Academy of Sciences 116:2612–2617.

Leisnham, P. T., L. P. Lounibos, G. F. O’Meara, and S. A. Juliano. 2009. Interpopulation divergence in competitive interactions of the mosquito Aedes albopictus. Ecology 90:2405–2413.

Levin, S. A. 1976. Population Dynamic Models in Heterogeneous Environments. Annual Review of Ecology and Systematics 7:287–310.

Mestre, A., R. Poulin, and J. Hortal. 2020. A niche perspective on the range expansion of symbionts. Biological Reviews 95:491–516.

Moran, E. V., F. Hartig, and D. M. Bell. 2016. Intraspecific trait variation across scales: implications for understanding global change responses. Global Change Biology 22:137–150.

Paine, R. T. 1966. Food Web Complexity and Species Diversity. The American Naturalist 100:65–75.

Pimm, S. L., and M. L. Rosenzweig. 1981. Competitors and Habitat Use. Oikos 37:1–6.

Post, D. M., E. P. Palkovacs, E. G. Schielke, and S. I. Dodson. 2008. Intraspecific variation in a predator affects community structure and cascading trophic interactions. Ecology 89:2019–2032.

Raerinne, J. 2020. Ghosts of Competition and Predation Past: Why Ecologists Value Negative Over Positive Interactions. The Bulletin of the Ecological Society of America 101:e01766.

Ricklefs, R. E. 2010. Evolutionary diversification, coevolution between populations and their antagonists, and the filling of niche space. Proceedings of the National Academy of Sciences 107:1265–1272.

Roughgarden, J. 1972. Evolution of niche width. American Naturalist:683–718.

Roughgarden, J. 1983. Competition and Theory in Community Ecology. The American Naturalist 122:583–601.

Schoener, T. W. 1983. Field Experiments on Interspecific Competition. The American Naturalist 122:240–285.

Soberón, J. 2007. Grinnellian and Eltonian niches and geographic distributions of species. Ecology Letters 10:1115–1123.

Soberón, J., and M. Nakamura. 2009. Niches and distributional areas: Concepts, methods, and assumptions. Proceedings of the National Academy of Sciences 106:19644–19650.

Tamme, R., I. Hiiesalu, L. Laanisto, R. Szava-Kovats, and M. Pärtel. 2010. Environmental heterogeneity, species diversity and co-existence at different spatial scales. Journal of Vegetation Science 21:796–801.

Thompson, J. N. 1988. Variation in interspecific interactions. Annual Review of Ecology and Systematics 19:65–87.

Tilman, D. 1982. Resource competition and community structure. Princeton university press.

Tilman, D. 2020. Plant Strategies and the Dynamics and Structure of Plant Communities.(MPB-26), Volume 26. Princeton University Press.

Travis, J. 1996. The Significance of Geographical Variation in Species Interactions. The American Naturalist 148:S1–S8.

Violle, C., B. J. Enquist, B. J. McGill, L. Jiang, C. H. Albert, C. Hulshof, V. Jung, and J. Messier. 2012. The return of the variance: intraspecific variability in community ecology. Trends in Ecology & Evolution 27:244–252.

Zélé, F., I. Santos, I. Olivieri, M. Weill, O. Duron, and S. Magalhães. 2018. Endosymbiont diversity and prevalence in herbivorous spider mite populations in South-Western Europe. FEMS Microbiology Ecology 94:fiy015.

