## Supplementary Material for "Shaken, not shifted: Genotypic variation tunes how interspecific competition shapes niches"

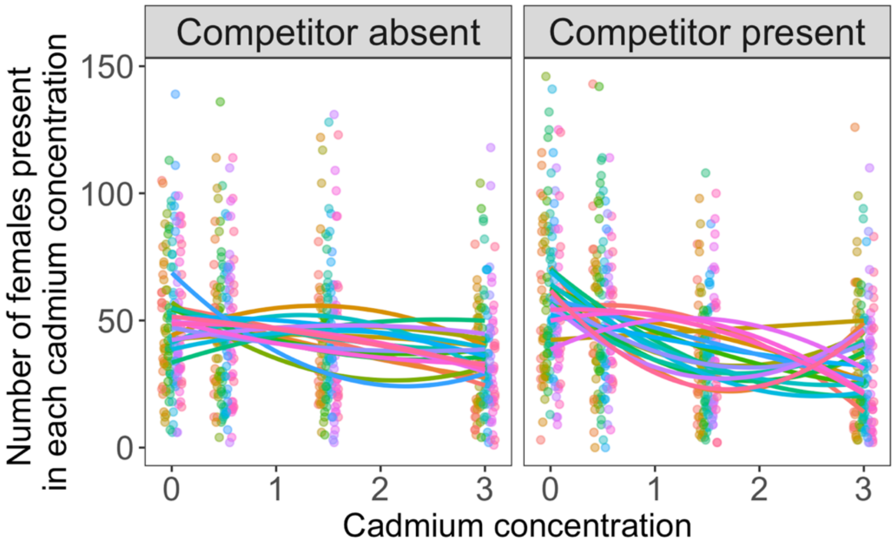


**Figure S1.** Variation in choice of cadmium concentrations between the 20 lines analyzed. The first panel corresponds to the treatment where there is a presence of the competitor on the two lowest cadmium concentrations and the second panel to the treatment without competitor.

**Table S1.** Polynomial model outcomes for the effects of interspecific competition on the number of females of *T. evansi* along a gradient of cadmium concentration.

| Line | Competitor absent | | | | Competitor present | | | | Change in curvature? |
|---|---|---|---|---|---|---|---|---|---|
| | First degree | Second degree | Intercept | Sign | First degree | Second degree | Intercept | Sign | |
| 4 | -0.09 | -0.06 | -1.15 | downward | 0.29 | -0.24 | -1.19 | downward | No |
| 5 | -0.24 | 0.00 | -1.13 | upward | -1.26 | 0.38 | -0.96 | upward | No |
| 8 | 0.34 | -0.12 | -1.46 | downward | -0.04 | -0.07 | -1.18 | downward | No |
| **9** | **0.07** | **-0.03** | **-1.38** | **downward** | **0.01** | **0.03** | **-1.39** | **upward** | **Yes** |
| **11** | **0.17** | **-0.05** | **-1.44** | **downward** | **-0.70** | **0.20** | **-1.11** | **upward** | **Yes** |
| 14 | -0.68 | 0.16 | -1.03 | upward | -0.82 | 0.21 | -1.02 | upward | No |
| 16 | -0.35 | 0.08 | -1.18 | upward | -0.38 | 0.02 | -1.03 | upward | No |
| 19 | -0.35 | 0.07 | -1.16 | upward | -0.87 | 0.23 | -1.00 | upward | No |
| **23** | **0.34** | **-0.07** | **-1.63** | **downward** | **-0.60** | **0.06** | **-0.92** | **upward** | **Yes** |
| 26 | -0.19 | 0.02 | -1.22 | upward | -0.75 | 0.21 | -1.09 | upward | No |
| **27** | **0.40** | **-0.17** | **-1.42** | **downward** | **-0.59** | **0.09** | **-0.98** | **upward** | **Yes** |
| **29** | **0.22** | **-0.07** | **-1.46** | **downward** | **-0.76** | **0.12** | **-0.89** | **upward** | **Yes** |
| **32** | **0.04** | **-0.05** | **-1.30** | **downward** | **-0.19** | **0.01** | **-1.19** | **upward** | **Yes** |
| **34** | **-0.80** | **0.17** | **-0.94** | **upward** | **-0.19** | **-0.03** | **-1.12** | **downward** | **Yes** |
| 38 | -0.19 | 0.04 | -1.27 | upward | -0.66 | 0.17 | -1.09 | upward | No |
| **42** | **0.15** | **-0.04** | **-1.45** | **downward** | **-0.90** | **0.28** | **-1.09** | **upward** | **Yes** |
| 44 | 0.01 | -0.06 | -1.25 | downward | 0.44 | -0.17 | -1.46 | downward | No |
| **51** | **-0.21** | **0.03** | **-1.20** | **upward** | **0.26** | **-0.18** | **-1.26** | **downward** | **Yes** |
| 54 | 0.04 | -0.06 | -1.27 | downward | 0.12 | -0.14 | -1.19 | downward | No |
| **55** | **-0.07** | **-0.03** | **-1.23** | **downward** | **-1.15** | **0.35** | **-1.00** | **upward** | **Yes** |

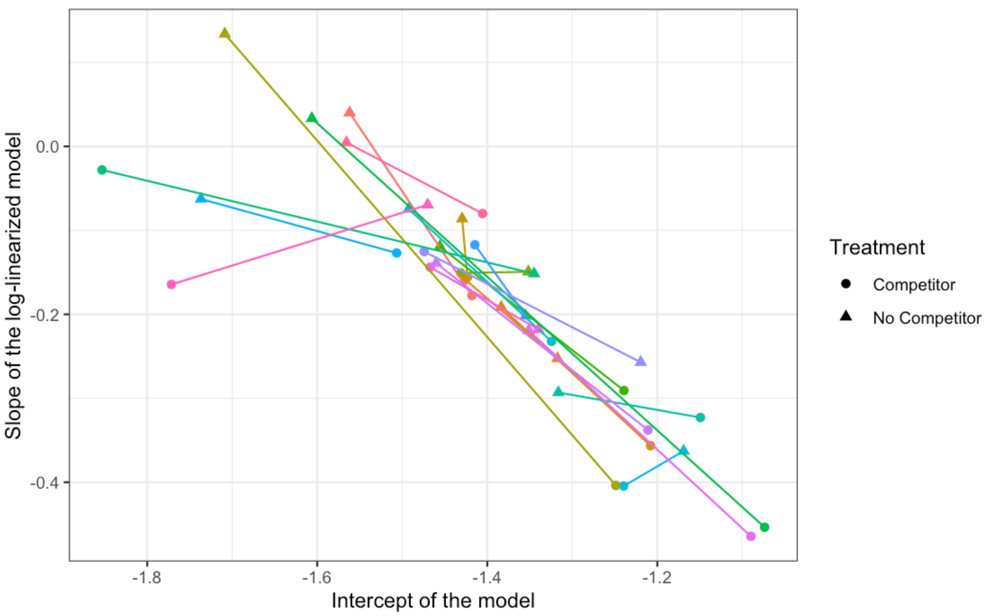


**Figure S2.** Relationship between the slope and intercept of log-linearized models to test habitat preference along the gradient of cadmium (i.e., log-transformed number of females as a function of cadmium concentration). Lines connect the responses of the same line in the “no competitor” (△) and competitor (O) treatments. No competitor results (△) show that most genetic lines have a negative slope (which indicates small decrease in preference with increased cadmium concentrations), while a few lines have almost no variation in abundance with cadmium concentration, and one line has a slight increase in performance with high cadmium concentration. Contrasting no competitor with competitor treatments (O), most genetic lines have a decrease in the slope (i.e., an increase in the choice towards the lower cadmium concentrations), but there are a few lines where the presence of the competitor increases the slope.

**Table S2.** Polynomial model outcomes for the effects of interspecific competition on the number of offspring per female of *T. evansi* along a gradient of cadmium concentration.

| Line | Competitor absent | | | | Competitor present | | | | Change in curvature? |
|---|---|---|---|---|---|---|---|---|---|
| | First degree | Second degree | Intercept | Sign | First degree | Second degree | Intercept | Sign | |
| **4** | **-0.23** | **0.42** | **0.23** | **upward** | **0.29** | **-0.24** | **-1.19** | **downward** | **Yes** |
| 5 | 0.01 | 0.45 | 0.30 | upward | -1.26 | 0.38 | -0.96 | upward | No |
| **8** | **-0.13** | **0.51** | **0.29** | **upward** | **-0.04** | **-0.07** | **-1.18** | **downward** | **Yes** |
| 9 | -0.25 | 0.28 | 0.22 | upward | 0.01 | 0.03 | -1.39 | upward | No |
| 11 | -0.62 | 0.79 | 0.48 | upward | -0.70 | 0.20 | -1.11 | upward | No |
| 14 | -0.38 | 0.95 | 0.31 | upward | -0.82 | 0.21 | -1.02 | upward | No |
| 16 | -0.10 | 0.24 | 0.36 | upward | -0.38 | 0.02 | -1.03 | upward | No |
| 19 | -0.20 | 0.31 | 0.31 | upward | -0.87 | 0.23 | -1.00 | upward | No |
| 23 | -0.36 | 0.47 | 0.40 | upward | -0.60 | 0.06 | -0.92 | upward | No |
| 26 | -0.30 | 0.31 | 0.40 | upward | -0.75 | 0.21 | -1.09 | upward | No |
| 27 | -0.08 | 0.15 | 0.20 | upward | -0.59 | 0.09 | -0.98 | upward | No |
| 29 | -0.16 | 0.20 | 0.24 | upward | -0.76 | 0.12 | -0.89 | upward | No |
| 32 | -0.14 | 0.33 | 0.33 | upward | -0.19 | 0.01 | -1.19 | upward | No |
| **34** | **-0.18** | **0.15** | **0.26** | **upward** | **-0.19** | **-0.03** | **-1.12** | **downward** | **Yes** |
| 38 | 0.61 | 0.15 | 0.25 | upward | -0.66 | 0.17 | -1.09 | upward | No |
| 42 | -0.02 | 0.13 | 0.20 | upward | -0.90 | 0.28 | -1.09 | upward | No |
| **44** | **0.18** | **0.17** | **0.24** | **upward** | **0.44** | **-0.17** | **-1.46** | **downward** | **Yes** |
| **51** | **-0.12** | **0.21** | **0.26** | **upward** | **0.26** | **-0.18** | **-1.26** | **downward** | **Yes** |
| 54 | 0.01 | 0.07 | 0.25 | upward | 0.12 | -0.14 | -1.19 | downward | Yes |
| 55 | 0.39 | 0.06 | 0.37 | upward | -1.15 | 0.35 | -1.00 | upward | No |
